# Low virulence evolves as a bet-hedging strategy in fluctuating environment

**DOI:** 10.1101/034041

**Authors:** Anne Nguyen, Etienne Rajon, David Fouchet, Dominique Pontier, Jorge Eduardo Rabinovich, Sébastien Gourbiére, Frédéric Menu

## Abstract

The effect of fluctuating environmental conditions (*i.e*. environmental stochasticity) on the evolution of virulence has been broadly overlooked, presumably due to a lack of connection between the fields of evolutionary epidemiology and insect ecology. Practitioners of the latter have known for a long time that stochastic environmental variations can impact the population dynamics of many insects, some of which are vectors of infectious diseases. Here we investigate whether environmental stochasticity affecting a vector’s life history can have an indirect impact on the evolutionarily expected virulence of the parasite, using Chagas disease as an example. We model the evolution of virulence using the adaptive dynamics framework, showing that parasite virulence should decrease when the vector's dynamics randomly change in time. The decrease is even more pronounced when environmental variations are frequent and ample. This decrease in virulence can be viewed as a bet-hedging strategy: when a parasite is at a risk of not being transmitted (e.g. because vectors are scarce), its best option is to stay in the host longer – that is, to be less virulent. Lowering virulence is thus very similar to increasing iteroparity, a well-known risk-spreading strategy, and should be expected to evolve whenever parasite transmission varies randomly in time.

## 1. Introduction

Virulence is the decrease in host fitness due to parasitic infection [1–3]. A parasite’s virulence presumably increases with its reproduction rate inside the host, as does its transmission between hosts (but see [4]). As a parasite must multiply before being transmitted, some virulence is required to ensure parasites transmission [1, 5], and an increase in virulence should mechanistically cause an increase in transmission [6, 7]. This relationship between virulence and transmission has been questioned in the past [8, 9] but seemingly relies on empirical (reviewed in [2, 10]) and theoretical grounds [2, 11].

The so-called trade-off between virulence and transmission is one of the pillars of theoretical evolutionary epidemiology, whose shape is known to drive the evolution of virulence [12]. When one ignores the competition between parasite strains within a host, an intermediate level of virulence is expected to evolve: a low-virulence parasite will scarcely be transmitted and may fail to infect hosts fast enough to persist, whereas a high-virulence parasite will kill its host shortly after infection, thus limiting its circulation in the host population [13, 14]. The exact level of virulence expected to evolve depends on the precise shape of the trade-off between virulence and transmission.

This shape may depend, for instance, on how parasites are transmitted from one host to another; this transmission can either be direct, or indirect through other species called vectors. Vector-borne parasites are theoretically expected to evolve a higher virulence to their hosts than directly transmitted parasites, for two reasons [15]. First, they are able to spread over long distances, which attenuates the evolutionary consequences of local host populations going extinct [16]. Second, high virulence sometimes leads to a decrease in hosts’ activity [17], which may decrease direct transmission but should have a null or positive impact on vector transmission. Virulence in vector-borne pathogens is therefore likely to evolve under a specific virulence-transmission trade-off [10, 18–20]. Moreover, vectors play a major role in pathogens transmission, such that their population dynamics are linked. Many vectors are insects whose life histories are highly sensitive to environmental changes of both biotic and abiotic origins *[e.g*. 21, 22]. Part of these changes are unpredictable (stochastic), and can have a strong impact simultaneously on all individuals of the population. Their impact can even be amplified by population regulation through density-dependent processes, yielding partly cyclic or chaotic population dynamics [23–25]. The evolutionary dynamics of virulence could be affected by these variations of a vector’s abundance [26], but this phenomenon has been largely neglected to this day.

Here we study the impact of stochastic variations affecting a vector’s demography on the evolutionarily expected virulence, using a model parameterized for the flagellate protozoan *Trypanosoma cruzi* – which causes Chagas disease [see 27, 28 for reviews] – and one of its vectors, the triatomine bug *Triatoma infestans*. These insects likely face unpredictable fluctuations of their environment, for instance due to punctual eradication campaigns in domestic areas and random climatic events elsewhere [29] – these environmental variations are currently increasing in amplitude and frequency due to ongoing global change (see [30]).We use an adaptive dynamics (AD) framework [31, 32] to predict the evolutionarily expected virulence of the pathogen which, as we show, decreases as the environment becomes more variable – both in frequency and in amplitude. Therefore, stochastic and deterministic environments should favour the evolution of strains with different virulence. We discuss these results in the light of the existing theory of evolution of life history traits in a fluctuating environment, showing that low levels of virulence can be considered as bet-hedging strategy, similar to iteroparity. Following this analogy, we expect our results to extend beyond the biological system modelled here and even beyond vector-borne diseases: low virulence is generally expected to evolve in fluctuating environments.

## 2. Material and Methods

We set up a Susceptible – Infected (SI) vector-borne epidemiological model in continuous time, and we use an evolutionary epidemiology approach based on AD [33] to study the evolution of a pathogen’s virulence.

### 2.1. The eco-epidemiological model

The model describes the evolution of virulence in a population of *T. cruzi* infecting a population of *Triatoma infestans*, the main vector of *T. cruzi* [34, 35] and a population of human hosts. Hosts and vectors are either susceptible (*Hs*, *Vs*) or infected by the strain *i* of *T. cruzi* (*Hi* and *Vi*; figure 1). We make the assumption of infrequent (or weak) mutations, such that there are at most two parasite strains in competition. We investigate the invasion success of a ‘mutant’ parasite strain in a population of hosts and vectors partially infected by a ‘resident’ parasite strain. The spatial scale is not explicitly taken into account but parameters correspond to populations of vectors and humans living in a homogenous area composed of a few houses [36, 37], which is the typical domiciliary rural landscape encountered by *T. infestans*. The basic time unit is one year but we use a resolution time of 8 hours for ordinary differential equations. Table 1 describes all the parameters of the model, including the symbols used, their description and definition, and their values.

**Figure 1.**
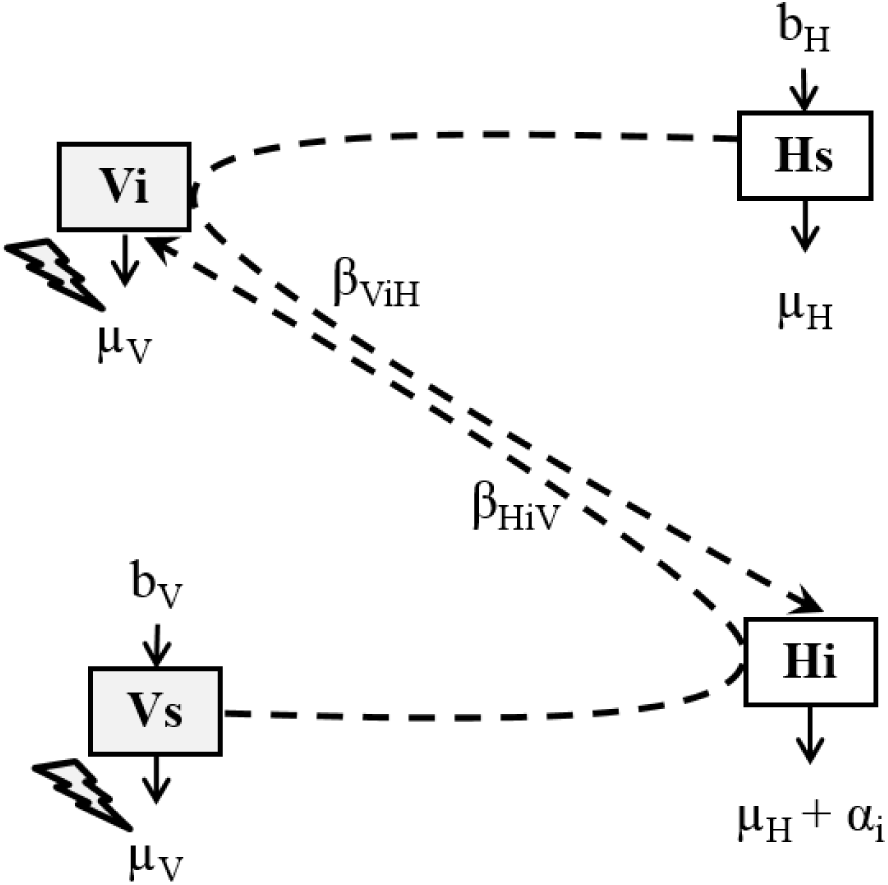
Schematic representation of the Susceptible – Infected model for one strain *i* of *T. cruzi*. Each compartment represents the number of individuals of interest: susceptible hosts (***Hs***), hosts infected by strain *i* (***Hi***), susceptible vectors (***Vs***) and vectors infected by strain *i* (***Vi***). Susceptible hosts become infected through contact with infected vectors according to the transmission rate *β_VH_* (dotted line). Susceptible vectors become infected through contact with infected hosts according to the transmission rate *β_VH_* (dotted line). We assume no vertical transmission so that susceptible and infected hosts and vectors give birth to susceptible offspring according to the rates *b_H_* and *b_V_* respectively (full line). Susceptible hosts die at constant rate *μ_H_* and infected hosts have an additive mortality *α* due to infection (full line). In the absence of environmental stochasticity (*i.e*. in deterministic environment) susceptible and infected vectors die at constant and same rate *μ_V_* (full line). Environmental stochasticity (lightning symbols) affects only vector natural mortality *μ_V_*, which thus varies between “good” and “bad” years independently of their infectious status. See Table 1 for further definitions of parameters.

**Table 1:**
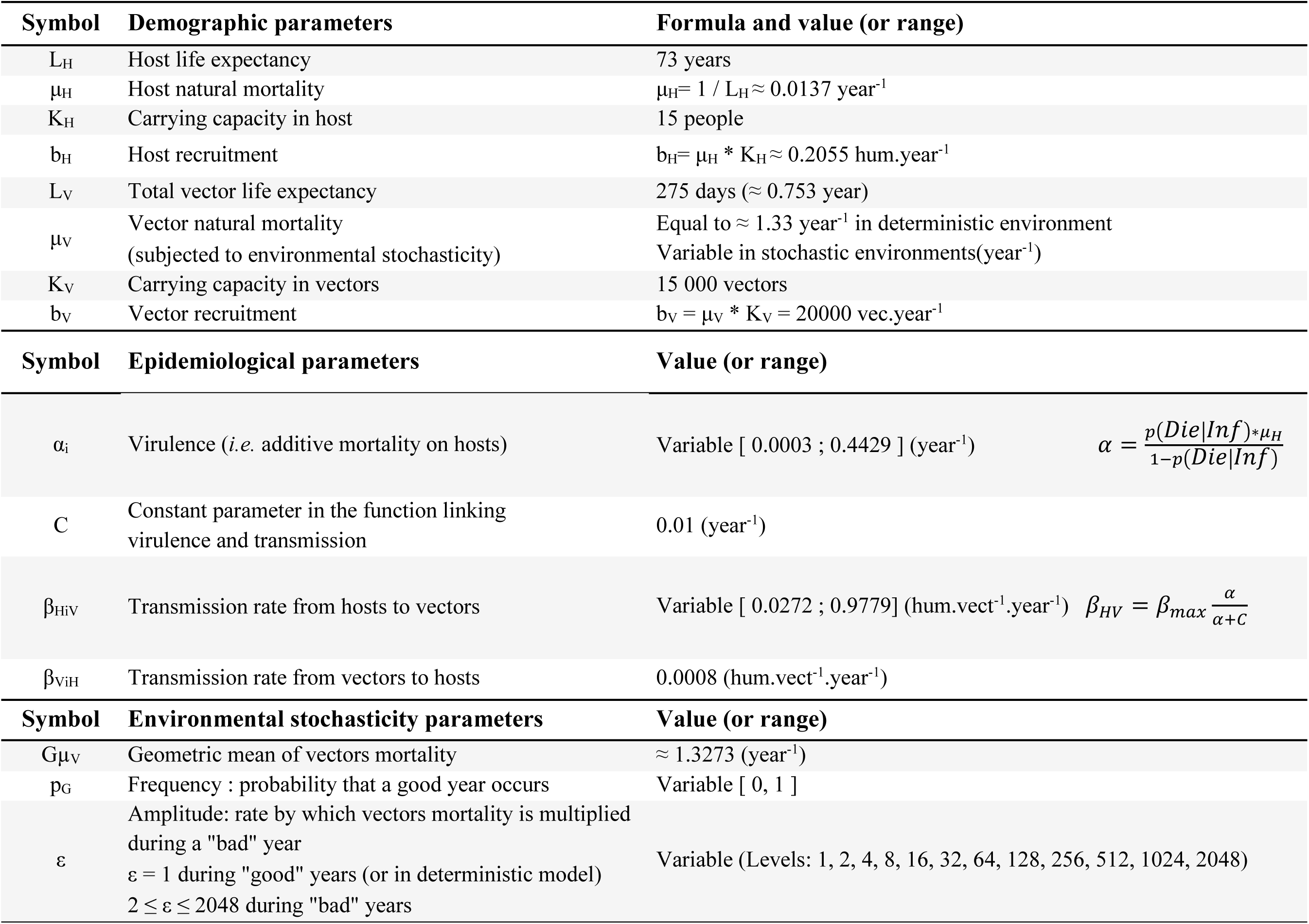
Model parameters: symbol, description, formula and value (or range). The first part presents hosts and vectors demographic parameters, the second the epidemiological parameters used in adaptive dynamics analysis and the third the environmental stochasticity parameters.

***Host and vector demography***. We calculate the hosts’ mortality *μ_H_* as the reciprocal of the average life expectancy *L_H_*. The basic information for the vector *T. infestans* was obtained from Rabinovich [38]. Hosts and vectors net recruitment rates (*b_H_* and *b_V_*) simultaneously account for birth and immigration and are considered constant. In the deterministic model with no parasite, the equilibrium host and vector population size (carrying capacity) are, respectively, 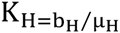 and 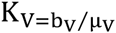.

The vectors’ mortality *μ_V_* depends on the environmental context that a population of vectors encounters, so it varies stochastically from one year to the next in the stochastic version of our model. We model environmental stochasticity as a random series of high and low values for the vectors’ mortality rate *μ_V_* [25, 26, 29]. Two parameters control the intensity of environmental stochasticity: the frequency of random events *p_G_* (probability that a good year occurs) and the amplitude of the variations in *μ_V_* denoted *ε* – precisely, *ε* is the ratio of bad-to-good vectors’ mortality:

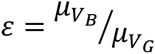 with *μ_VB_* and *μ_VG_* the vector mortality during bad and good years, respectively 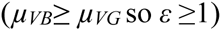.

Changing *μ_VB_* or *μ_VG_* would affect both the variance and mean of the vector mortality. In order to study the impact of environmental variability only, we kept the geometric mean of vector mortality *G_μv_* constant across simulations, equal to a value estimated under laboratory conditions. We use the definition of the geometric mean [25]:

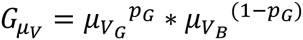 to calculate the values of *μ_VG_* and *μ_VB_* for each amplitude *ε* and frequency of random events *p_G_* of environmental stochasticity considered, which yields (see electronic supplementary material, appendix A for detailed calculation): 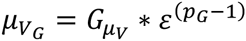 and 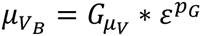

We also performed the analysis by keeping the arithmetic mean of vector mortality *A_μv_* constant across simulations. The qualitative trends are similar in the two cases (results not shown).

We consider various environmental contexts from deterministic (*ε* = 1: no difference in vector mortality between “good” and “bad” years) to highly stochastic (*ε* = 2048). The latter means that vector mortality during “bad” years is 2048 times higher than vector mortality during “good” years. Even though the amplitude of environmental stochasticity in life-history parameters is often largely unknown, the range of values of *ε* that we used should cover most plausible environments. We also study the impact of the frequency of random events expressed as the probability of occurrence of a good year *p_G_* (see Table 1). Note that if *p_G_* equals 0 or 1, the environment is deterministic for any value of *ε* – just as if *ε* equals 1.

***Parasite transmission***. We only consider vector transmission, which is the main mode of transmission for *T. cruzi* within its endemic geographic area. Thus, we ignore vertical transmission (from mother to offspring) and “accidental” transmission due to blood transfusion, organ transfer or contaminated food [see 39, 40, 41]. Susceptible hosts can become infected through a vector’s faeces laid close to a bite scar following a blood meal; the parasite may enter the body when hosts scratch themselves. Susceptible vectors may ingest parasites during blood meals on infected hosts and become infected. The transmission rate from infected vectors to hosts is denoted *β_ViH_*, and that from infected hosts to vectors is denoted *β_HiV_*. They result from the product of the biting rate and the probability of transmission – which do not appear explicitly in the model [see 42, 43]. At the population scale, the transmission from infected vectors to susceptible hosts is the product of *β_ViH_* (set to 0.0008 hum.vect^-1^.year^-1^), the density of infected vectors *Vi*, and the probability that the bitten host is susceptible – that is, the frequency of susceptible hosts *Hs/*(*Hs+Hi*). Similarly, the overall transmission rate from infected hosts to susceptible vectors is the product of the transmission rate *β_HV_*, the density of susceptible vectors *Vs* and the probability that the bitten host is infected *Hi*/(*Hs*+*Hi*). We assume no recovery, no co-infection (several strains in the same host or vector) and no super-infection (once infected, hosts or vectors cannot be infected by another strain).

*T. cruzi* strains differ by their levels of virulence *α_i_* and by their transmission rate *β_HiV_*. The virulence was defined in accordance with [4] and [44]:

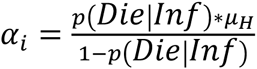 (1), where *p*(*Die\Inf*) is the probability for an individual host to die from the disease once infected. The total mortality rate of infected hosts is the sum of natural mortality and additive mortality: (*μ_H_+ α_i_*) year^-1^.

Virulence and transmission rates from host to vector are linked by a trade-off, as discussed earlier, such that high transmission only prevails when virulence is high:

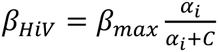 (2), where *β_max_* is the maximum transmission rate (set arbitrarily to 1 hum.vect^-1^.year^-1^). *C* defines the shape of the trade-off [32], which we set to 0.001 year^-1^ according to [26].

### 2.2. Evolutionary analysis with AD

The analytical analysis model is based on the calculation of the basic reproductive number *R_0_* [see 45, 46, 47 for reviews]. *R_0_* is a reliable estimate of a parasite’s fitness in a deterministic environment with simple population regulation, but it is inadequate when random variations are important determinants of the population dynamics – but see [48] and [49] for a derivation under seasonal fluctuations in a vector population. Studying evolution in a complex context involving both density dependent regulation of vectors’ populations and environmental variations imposes numerical simulations; we conducted them in both the stochastic and the deterministic context, the latter to validate our numerical model.

AD consists in studying the invasion success of a rare mutant strain that appears in host and vector populations already and partially infected by a resident strain [50]. It considers the fate of a rare mutant in competition with a high-frequency resident genotype – note that we use ‘genotype’ for simplicity, but any highly heritable medium of phenotypic information would apply. The success of the mutant at invading the resident’s population is evaluated through a metrics called the invasion fitness. The evolutionary outcome is then analysed by considering the outcome of the competition between every potential pairs of mutant and resident genotypes [31, 51–53].

#### 2.2.1. Analytical analysis of equilibrium in a deterministic environment

##### Model equilibrium and stability of the endemic equilibrium

In the analytical analysis, we first considered the system of transmission of *T. cruzi* with only the resident strain (sub-index r):

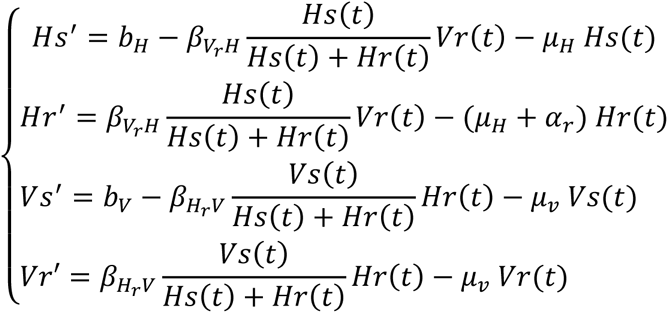

From this ODE system, we determined its equilibrium denoted {Hs^*^, Hr^*^, Vs^*^, Vr^*^}. We then introduced a mutant (sub-index *m*), with a frequency low enough that its impact on the dynamics of hosts and vectors can be neglected. At the time of invasion, the dynamics of the mutant and resident can be modelled as:

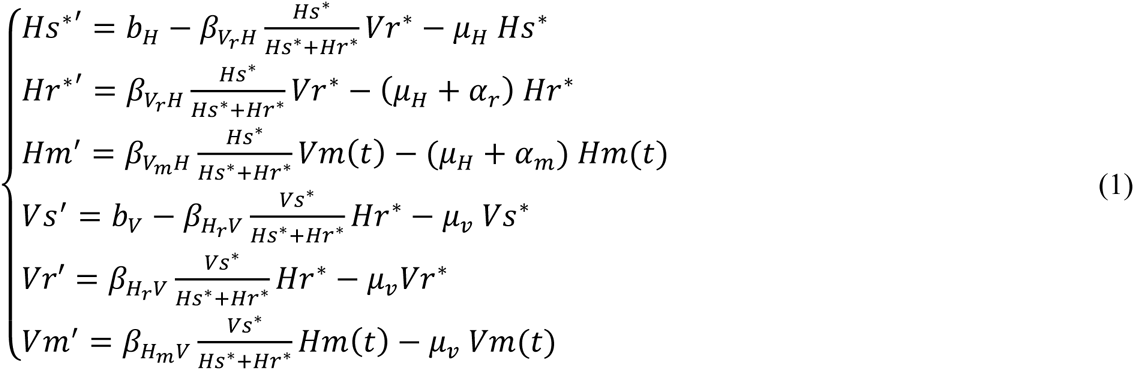

We determined the Jacobian matrix of the system with the two ordinary differential equations Hr’ and Vr’ [50]. We expressed the determinant and the trace of this matrix to analyse the stability of the equilibrium (see electronic supplementary material, appendix B).

#### 2.2.2. Numerical simulations in deterministic and stochastic environments

The analytical analysis performed above is out of reach in the presence of environmental stochasticity, so the invasion fitness needs to be estimated numerically *[e.g*. 25, 54]. We also performed numerical simulations in the deterministic context for validation. We assess the fate of a mutant strain (*α_m_*) emerging in a population of resident pathogen (*α_r_*) by simulating a competition between the two strains along a random path of “good” and “bad” years – where vector mortality is low or high, respectively. µ_v_ hence calculated is then used in a new EDO system derived from the system (1), such that the dynamics of the hosts and vectors are repeatedly (and stochastically) perturbed. The population dynamics was simulated with the resident population alone for 10 000 years – after checking that this time is largely sufficient for the resident population to reach the equilibrium in the deterministic environment. Then one mutant was introduced in the host and vector populations, spread between the hosts (frequency *HR*/(*HR*+*VR*)) and vectors (frequency *VR*/(*HR*+*VR*)). We simulated the resident’s and the mutant’s population dynamics during another 590 000 years – with no change in our results compared to simulations over 390 000 years.

We recorded the initial and the final densities of the mutant, and calculated the ratio of the final-to-initial density. This ratio is larger than 1 if the mutant tends to increase in number in the resident’s population. Otherwise – if the ratio is lower than 1 – the mutant eventually goes extinct. Because simulations run for a finite sequence of random events, a neutral mutant’s growth rate may sometimes deviate from the neutral expectation of 1. For this reason, we calculated the invasion fitness of a mutant as the difference between its own growth rate and that of the neutral mutant – *i.e*. that of the mutant with the same phenotype as the resident. A mutant with an invasion fitness larger than 0 will increase in frequency on average, and eventually invade; otherwise the mutant will go extinct.

We repeated this operation for every couple of resident and mutant strategies, using the exact same sequence of random events for all couples. We considered a large set of values of virulence *α*, between 3.10^−4^ and 0.4429 year^−1^, and for each, we obtained *β_HV_* from equation (2) (see Table 1). Then we plotted the areas where the invasion fitness is larger and lower than 0 on a Pairwise Invisibility Plot (PIP), for all couples of resident and mutant phenotypes in this set. This representation allows the identification of various types of evolutionary strategies [53]. We only obtained one type, Continuously Stable Strategies (CSS), which are both Evolutionarily Stable (ESS) – *i.e*. they cannot be invaded by any other strategy once established in the population – and Convergent-Stable – *i.e*. they can be reached through small mutational steps [see 33, 53]. Due to these two properties, these strategies are likely to stand as the ending state of adaptive evolution. We performed this analysis in 12 × 11 environmental contexts (*ε* = {1, 2, 4, 8, 16, 32, 64, 128, 256, 512, 1024, 2048} and *p_G_=* {0,0.1,0.2,…,1}), producing 132 Pairwise Invasibility Plots that we analyzed graphically to obtain the CSS. We restricted the range of virulence between 0.01120 and 0.01175, and checked that no other singular strategy exists within the range defined in Table 1. As shown in figures 2 and 3, we plotted a second PIP with a higher definition close to the ESS, in order to estimate more precisely the CSS.

**Figure 2.**
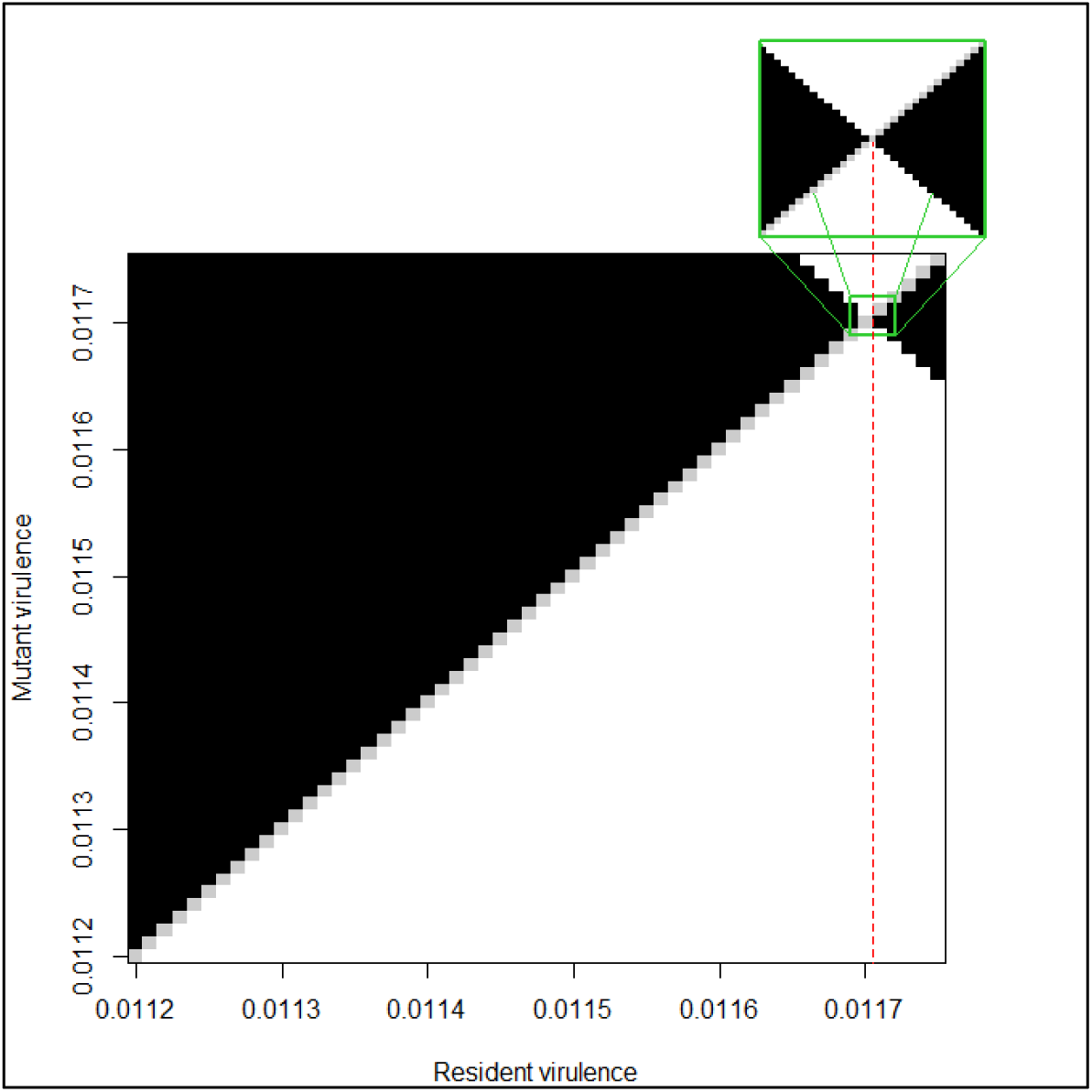
Evolutionary analysis with a Pairwise Invasibility Plot (PIP) in deterministic environment (ε =1). For all pairs of resident (*α*_R_ - x axis) and mutant parasites (α_M_ - y axis) the outcome of the competition is shown as black (the mutant invasion fitness increases above 1.10^−8^), white (the mutant invasion fitness decreases below −1.10^−8^) or grey (neutral mutant invasion fitness is between −1.10^−8^ and 1.10^−8^). The intersection of the lines separating black and white areas indicates the Continuously Stable Strategy (CSS) that cannot be invaded by any other strategy once established in the population and can be reached via small mutational steps. The principal PIP represents virulence between 0.0112 and 0.0117 with a resolution of 10^−5^. Around the CSS revealed, we used a resolution of 10^−6^to reach the CSS with a greater precision, which is represented in the little PIP above, intentionally enlarged. In this deterministic case, the CSS corresponds to the strain with a virulence α^*^ of 0.011705.

**Figure 3.**
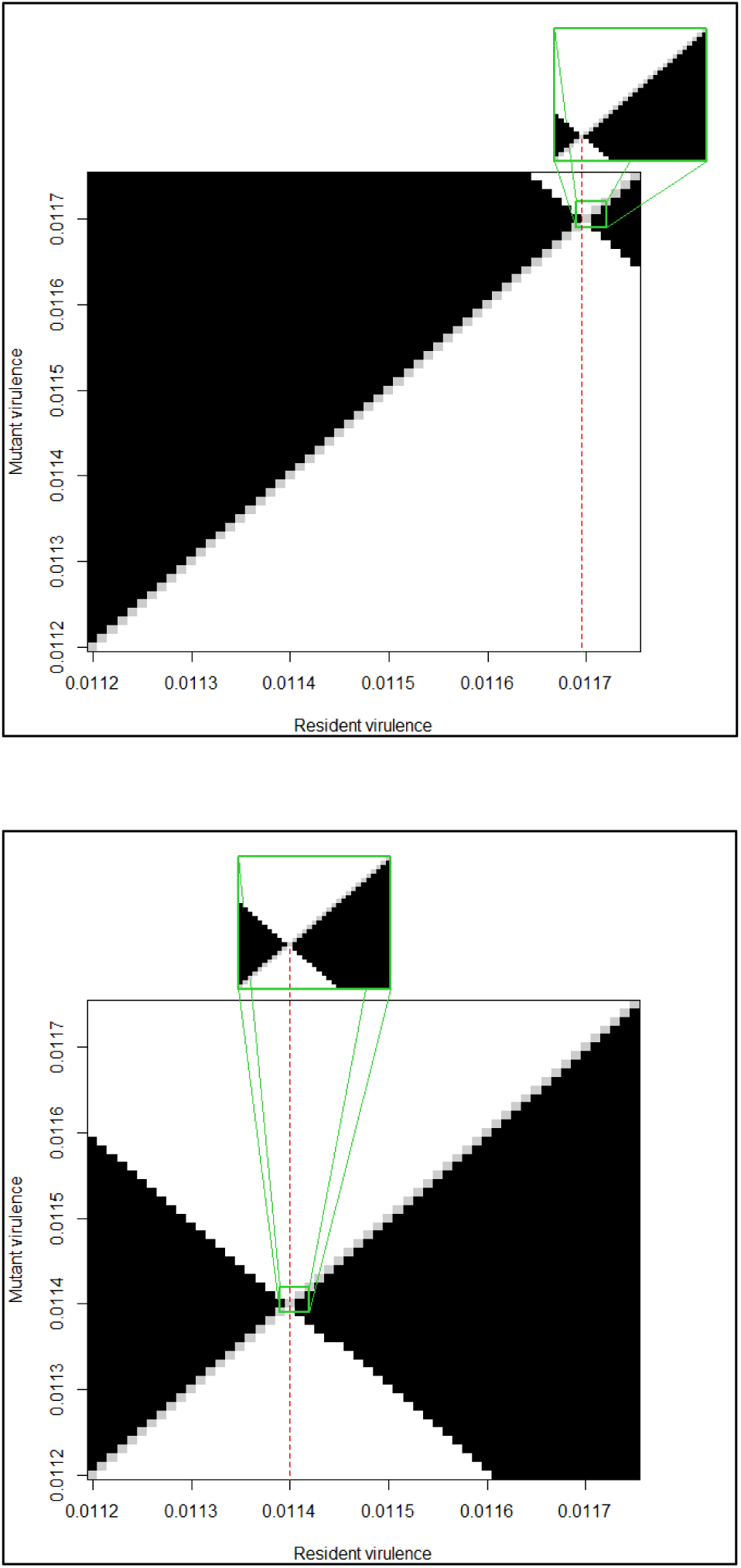
The evolutionarily expected virulence decreases as the environment becomes more variable. In both PIPs, there is a unique singular strategy, which is a CSS. In the low stochastic environment (ε = 2, *p_G_* = 0.5, panel A), the CSS corresponds to a strain with a virulence *α^*^* of 0.011695, whereas in the more variable environment (*α* = 128, *p_G_*= 0.5, panel B) we expect a lower virulence *α*^*^ of 0.0114 to evolve.

## 3. Results

### 3.1. The evolution of virulence in a deterministic environment

#### 3.1.1. Analytical analysis: infering the condition under which a mutant strain can invade the endemic state

In the deterministic version of our model, the epidemiological rates (*β_vHr_, β_HVr_, α_r_* and *β_VHm_, β_HVm_, α_m_*) are constant and density independent, a convex trade-off is considered between virulence *α* and transmission from hosts to vector *β_HV_* and no co-infection or superinfection is allowed. Therefore, as for directly transmitted disease [55], an optimal combination of parasite traits is expected to evolve, which maximizes the *R_0_* of the mutant [32]. In our study, we find that the strain with the highest *R_0_* (12.8) has a virulence *α*^*^ of 0.011704.

#### 3.1.2. Numerical simulations

The results of the numerical simulation are represented as Pairwise Invasibility Plots (figure 2). Each pair of resident parasites (*α_R_ - x axis*) and mutant (*α_M_ - y axis*) are represented as a square, whose colour represents the result of mutant invasion – black for successful invasion (mutant growth rate above 10^−8^), grey for quasi-neutrality (growth rate between −10^−8^ and 10^−8^), and white for failure to invade (growth rate below −10^−8^). We find a singular strategy at the intersection of the two lines separating black and white areas. This strategy is Continuously Stable, *i.e*. it can be reached by small mutational steps and, once established in the population, cannot be invaded by any other strategy. The Continuously Stable Strategy, at *α*^*^ = 0.011705, is similar to the virulence expected from the analytical approach using *R_0_*.

### 3.2. The evolution of virulence in stochastic environments

We first consider an environment where *p_G_* = 0.5, such that vector’s survival is equally likely to be high or low (figure 3). We study the evolution in two markedly different contexts: one where the amplitude of variation in vector mortality *μ_V_* is low (*ε* = 2, figure 3 A), the other where it is high (*ε* = 128, figure 3 B).In the first context, we find that the evolutionarily expected virulence *α^*^* equals 0.011695, lower but close to that found in a deterministic environment (0.011705). However, when variations are more pronounced (*ε* = 128), the selected virulence *α^*^* decreases drastically to 0.0114.

A more thorough analysis of the impact of the amplitude of environmental variations *ε* and of the frequency of random events *p_G_* is presented in figure 4. The relation linking the evolutionarily expected virulence *α^*^* and *p_G_* is non-monotonic: *α^*^* first decreases as *p_G_* increases towards 0.4 to 0.6 – depending on the amplitude of variation *ε* – and then increases as *p_G_* approaches 1. This non-monotonicity can be explained by the fact that the number of changes – from a good to a bad year or the opposite – is minimum when *p_G_* is either small or large. The evolutionarily expected level of virulence *α^*^*also decreases as the amplitude of variation, ε, increases. This result holds as long as *p_G_* differs from 0 or 1 (*i.e*. deterministic environment).

**Figure 4.**
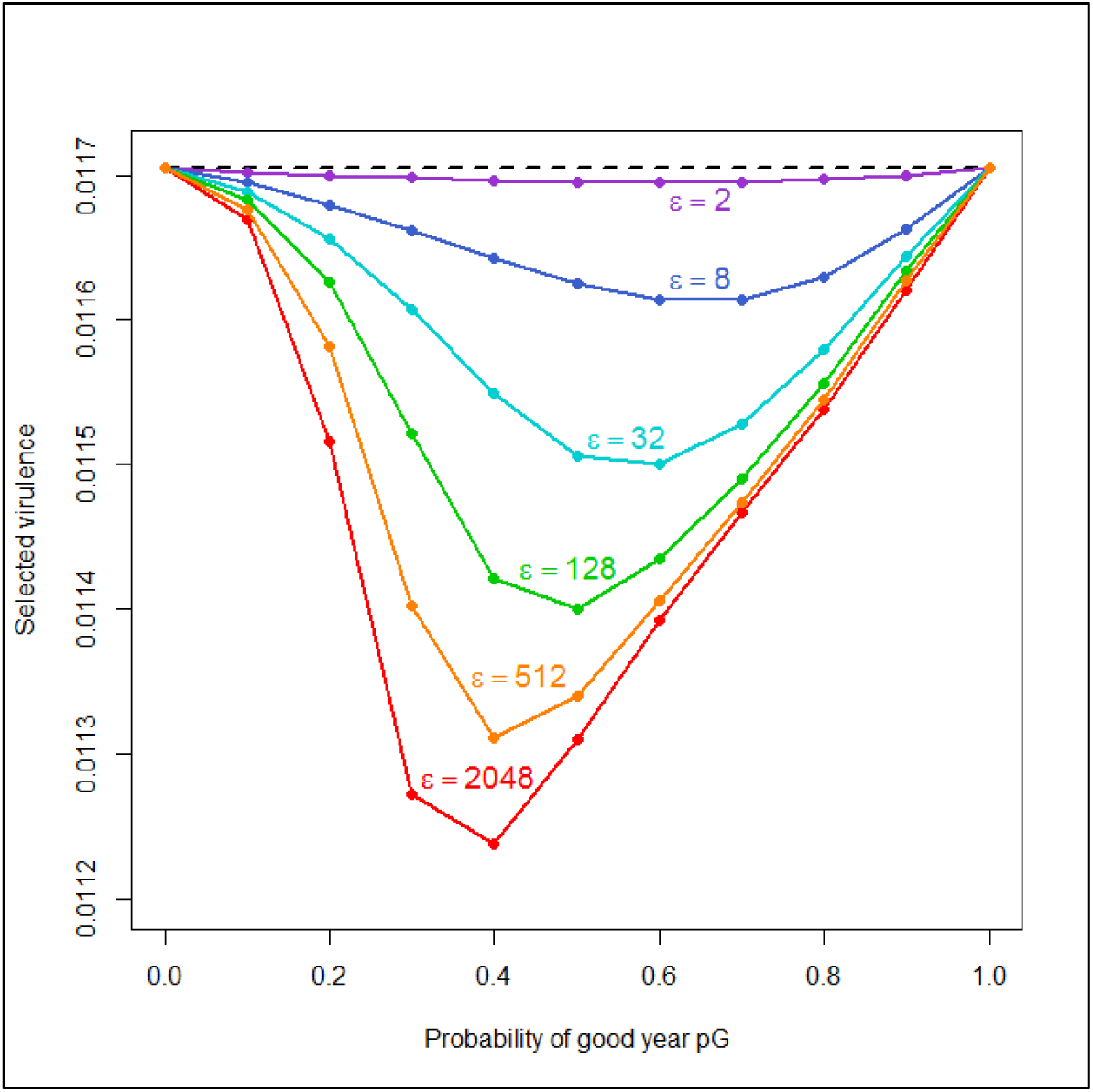
Influence of environmental stochasticity parameters *ε* and *p_G_* on virulence evolution. The graph represents the evolutionarily expected selected virulence (*α^*^*) as a function of the probability that a good year occurs *p_G_*, for different intensities of environmental stochasticity ε. The selected virulence decreases with an increase in the probability of occurrence of a good year *p_G_* until a threshold is reached and increases afterwards. The decrease of the selected virulence and the thresholds are greater as the amplitude of environmental stochasticity *ε* is higher.

## 4. Discussion

Using a standard evolutionary epidemiology analysis (*R_0_*) and simulations, we find the evolutionarily expected level of virulence in a stable and predictable (deterministic) environment, which results from a trade-off between host survival and parasite transmission. We show that the virulence expected to evolve in a stochastic environment is lower than that expected in a deterministic context, with a difference that increases as environmental variations become more frequent and wider in amplitude.

### Why would a parasite lowering its virulence be selectively favoured in an unpredictable environment?

Consider first a highly virulent parasite that would kill its host shortly after infection – ignoring that this would often be sub-optimal even in a stable environment. This parasite would likely kill its host before the next environmental change. Transmission needs to occur during the host’s lifetime, such that any parasite in the population will either encounter a high or a low density of vectors – and thus have a high or a low transmission rate – during its time within a host. Therefore, the transmission of this strain would be very variable over a long time period, and more so if environmental changes are frequent or have a pronounced impact on vector densities. In the long term, the number of infected hosts would decrease because of this high variance in transmission rate, just as a population growth rate decreases when survival or fecundity vary in time [56].

Conversely, a lower virulence extends the lifetime of the infected host and of its low-virulent parasite, which may thus encounter various environmental conditions (*i.e*. various vector densities). This reduces the temporal variance in the transmission of the parasite, because its overall transmission rate is the sum of the rates encountered during the host’s lifetime. As an illustration, consider a parasite that lives two years, in a yearly-changing environment that can either contain a low (L) or a high density (H) of vectors, each with probability 0.5. Its probability of experiencing two L-years – and to have the lowest possible transmission rate – is 0.25, the same as two H-years – and the highest possible transmission rate. The remaining possible situation is one L- and one H-year, with a transmission rate equal to the lineage average over an infinite time. This parasite thus has a reduced variance in transmission rates compared to a parasite that would kill its host within a year, which would have either the lowest or the highest possible transmission rate. As the lifetime of the host increases, combinations that yield extreme transmission rates become less likely, such that temporal variance in the transmission rates decreases.

This decrease in virulence in order to buffer the temporal variance of transmission rates can thus be viewed as bet-hedging at the level of the parasite. Actually, it is nearly identical to a well-known bet-hedging strategy, the iteroparity. Iteroparous individuals reproduce several times, and their longevity depends on adult survival (similar to virulence in our model). An increase in longevity provides more opportunities for reproduction, which may increase or decrease the overall lifetime reproductive success depending on the shape of a trade-off between litter size (somehow similar to the transmission rate in our model) and adult survival [57]. In a deterministic environment, the shape of the trade-off is generally sufficient to predict the evolutionarily expected position on the trade-off; that is, the evolutionarily expected litter size and adult survival. In a fluctuating environment, however, selection may favour an increase in adult survival, as it provides more reproduction opportunities and thus reduces the temporal variance in the lifetime reproductive success [58], which is the definition of a bet-hedging strategy [59].

The analogy between iteroparity and virulence is rather straightforward: a decrease in virulence allows the parasite to live longer within its host, such that it has more occasions of being transmitted. In a fluctuating environment, this reduces the variance in its success at being transmitted and, in the long term, it will infect more hosts. We considered environmental factors affecting the vector’s dynamics, but by analogy with other bet-hedging strategies we can presume that chaotic vector dynamics may also select for a decrease in virulence [24]. Furthermore, this analogy between iteroparity and virulence extends our results beyond the specific case of vector-borne diseases: any source of random variation that would impact transmission in the whole population simultaneously should select for lower virulence.

The relative impact of environmental stochasticity should depend on other selection pressures acting on virulence, such as the shape of the trade-off between virulence and transmission [18]. Environmental stochasticity should only play a minor role in the presence of a trade-off that selects strongly for a specific virulence in a stable environment, whereas it should significantly decrease the evolutionarily expected virulence whenever the trade-off conveys weak selection on virulence.

These theoretical developments into the evolution of virulence might allow us to explain some of the existing diversity in virulence among pathogens. For instance, *T. cruzi* is thought to be a highly genetically diverse parasite [60], commonly partitioned into six discrete typing units (DTUs TcI – TcVI) with several genotypes described in each DTU [61]. This high diversity might be partly explained by different environmental contexts, and the results of our study suggest that environmental variance might play an important role, especially in a context where transmission is spatially restricted and where triatomine species and trypanosome genotypes are non-randomly associated {Vallejo, 2009 #143}.

Differences in the life-history strategies of the vectors provide another possible explanation for this diversity. Vectors could also evolve bet-hedging strategies, such as facultative prolonged dormancy known in many insects [e.g. 29, 62] and iteroparity -which is observed among the Triatomines at variable level. By doing so, they reduce the temporal variance in their population size, and thus the temporal variance in the transmission rate of the parasite. This would decrease the selection pressure on the parasite for a bet-hedging strategy – *i.e*. for a decrease in virulence, such that the evolving virulence should depend on both the environmental context and the population and evolutionary dynamics of the vector. This interaction between the vector’s and parasite’s evolutionary dynamics was suggested by [26], who found that more virulent strains might invade at short timescale when the vector buffers environmental variance *via* a risk-spreading strategy. Nevertheless, the long-term coevolution of vectors and parasites’ traits in a fluctuating environment should be studied more thoroughly, as it might play a key role in the evolution of virulence.

## 5. Acknowledgments

This work has been supported by the French National Research Agency (grant reference “ANR-08-MIE-PP7”) and by the Agencia Nacional de Promoción Científica y Tecnológica of Argentina (grant reference PICT2008-0035), and benefited from an “Investissement d’Avenir” grant managed by the French National Research Agency (CEBA, ANR-10-LABX-25-01). This work was performed within the framework of the LABEX ECOFECT (ANR-11-LABX-0048) of University of Lyon, within the program “Investissements d’Avenir” (ANR-11-IDEX-0007) operated by the French National Research Agency (ANR).We are very grateful to V. Miele, M. Ginoux and A. Siberchicot for their kind assistance with the programming language and to S. Alizon for his comments on a previous version of the manuscript. This work was performed using the computing facilities of the CC LBBE/PRABI.

## 6. Competing interest

We have no competing interest.

## 7. Authors contributions

FM and AN designed the study. AN built the model, supervised by FM and JR. DF, ER and SG checked and improved the model. JR produced biological values of vector demographic parameters. AN analyzed the results and ER, DF, DP, SG and FM contributed to their interpretation. AN and ER wrote the manuscript with contributions from SG and FM. DF, DP, and JR revised the manuscript critically. All authors approved the final version of the article.

